# Effect of tempo on newborns’ neural processing of auditory rhythm: Emergence of sensitivity to metrical structure

**DOI:** 10.64898/2026.07.07.736947

**Authors:** Mohammadreza Edalati, Maya Psaris, Alban Gallard, Arthur Foulon, Fabrice Wallois, Barbara Tillmann, Laurel J. Trainor, Sahar Moghimi

## Abstract

Rhythm perception ability underpins music and language processing, and its developmental origins trace back to the earliest periods of life. While most rhythm patterns contain a variety of inter-onset intervals, listeners typically extract a steady underlying beat periodicity, as well as beat grouping periodicities (groups of two or three beats will be at frequencies 1/2 or 1/3, respectively, of that of the primary beat), forming a metrical hierarchy. While the developing brain can track auditory rhythm before birth, a previous study found that neural sensitivity to faster beat-related periodicities emerges early in third trimester of gestation, whereas encoding of slower metrical structure periodicities only appears closer to term birth. However, as rhythm patterns were only presented at one tempo, neural sensitivity to metrical structure could not be disentangled from sensitivity to tempo during early development. Thus, here we presented auditory rhythmic sequences at two different tempi and used high-resolution electroencephalography to measure neural sensitivity to their encoding in full-term newborns and young adults. Adults demonstrated a similar sensitivity to metrical structure across tempi, with greatest response at the duple metrical frequency regardless of tempo. Newborn neural responses, by contrast, were tempo-dependent, displaying markedly different response patterns across beat and meter frequencies at the different tempi. Together, these results reveal that tempo and metrical structure interact in shaping how the neonatal brain encodes auditory rhythm, suggesting that early neural processing may be constrained in its ability to track slower periodicities, and highlighting a developmental shift in the relative contributions of tempo and metrical structure to rhythmic processing.

**Research Highlights:**

- Adults show stable sensitivity to metrical structure across tempi, with peak responses at the duple level.
- Newborns’ neural responses are tempo-dependent, showing distinct patterns across beat and metrical frequencies.
- Tempo and metrical structure jointly shape neural encoding of rhythm in the brain, revealing a developmental shift in their relative contributions.

## Introduction

Rhythm is a fundamental organizing principle of both music and language, and plays a critical role in early development (Choi et al., 2020; Cirelli et al., 2018; Lense et al., 2021; Savage et al., 2021; Snyder et al., 2024; Tillmann et al., 2025). From birth, infants are immersed in rich rhythmic environments, where temporal regularities structure early infant-directed communication (Alviar et al., 2026). While rhythm patterns typically contain event inter-onset intervals of different durations, the brain extracts a regular underlying beat (what people tap to) from this rhythmic surface. Further, rhythmic structure in both music and language is inherently hierarchical. In music in particular, beats at one level are grouped (typically in groups of two or three) to create temporal regularities at multiple slower nested levels, together forming a metrical hierarchy (Fiveash et al., 2021; Large et al., 2023) that unfolds over distinct temporal scales or frequencies (Goswami, 2022; Trainor & Marsh-Rollo, 2018). Successful processing of rhythmic structures requires extracting and encoding these multiple hierarchical levels. While acoustic energy at hierarchical levels is typically present in a rhythmic stimulus, adults encode and perceive metrical levels even when there is little energy in the stimulus at those frequencies (Nozaradan et al., 2012), suggesting that metrical perception involves cognitive processing and expectancy-based abstraction. Developmental studies have shown that perception of metrical structure develops progressively across infancy and childhood, and is strongly shaped by experience (Hannon & Trainor, 2007; K. Nave et al., 2024; K. M. Nave et al., 2021). Recent studies provide evidence for early neural encoding of auditory rhythmic hierarchies (Edalati et al., 2023; Háden et al., 2015; Lenc et al., 2023). However, it remains unclear how dependent this might be on tempo. If infants are less able to track slower frequencies, their ability to track metrical levels at slower tempi could be impeded. Alternatively, infants might have a more narrow tempo range over which they can track metrical structure in general. As the relative contributions of frequency and metrical structure in shaping early neural responses to rhythm remain poorly understood, we explore this question here.

By the middle of the first year after birth, electroencephalographic (EEG) studies have revealed infants have some sophisticated meter perception abilities. Around 5-6 months, they can extract metrical structures from rhythm patterns even when there is low acoustic energy at the corresponding meter frequencies (Lenc et al., 2023). Further, presenting 6-month-olds with an ambiguous rhythm that can be perceived as either in duple or triple meter reveals that they can be biased via auditory priming to better encode one meter over the other (Flaten et al., 2022). However, the range of tempi over which such meter perception operates in early development remains unknown. Between 6 and 12 months, infants become specialized for processing the metrical structures dominant in the music in their environment (Hannon & Trehub, 2005b; Soley & Hannon, 2010), similarly to how they become specialized for the speech sounds in the language of their culture during this time (Kuhl, 2011; Werker & Tees, 2005). Beat and meter perception continue to develop across school years (K. Nave et al., 2024), with adult-like meter perception emerging in young adulthood (Nave-Blodgett et al., 2021a, 2021b).

Near-term and full-term infant brains encode auditory rhythms (Bianco et al., 2025; Edalati et al., 2023; Mashhadi et al., 2025; Winkler et al., 2009) and even show evidence for predictive timing of the underlying beat (Edalati et al., 2024; Háden et al., 2024). Saadatmehr et al. (2025) recently demonstrated that while the beat frequency of a presented rhythm pattern was neurally encoded by the beginning of the third trimester, the slower periodicities corresponding to the metrical structure were not robustly encoded until near the age of term. However, as the rhythm was presented at only one tempo, this raises the question of whether the youngest infants were unable to encode metrical structure, or whether they were unable to track slow periodicities. Yet, understanding this remains essential for clarifying the early developmental mechanisms by which the brain organizes and abstracts temporal information in music and language.

To disentangle the respective influence of metrical structure and tempo, we conducted an EEG study in adults and newborns using the 6-beat rhythm first used by Phillips-Silver & Trainor (2005) that has acoustic energy at both beat and nested metrical frequency levels. This repeating six-beat rhythm is ambiguous in containing evidence for both two-beat (three groups of two beats, hereafter referred to as duple meter) and three-beat (two groups of three beats, hereafter triple meter) groupings (see Fig. 1A). In Saadatmehr et al. (2025), the rhythm was presented at only one tempo, with the duple meter frequency being half that of the beat frequency, and the triple meter frequency one-third that of the beat frequency. They found that robust neural responses to the slower metrical periodicities only emerged near the age of term (as was also observed in Edalati et al. (2023) using the same pattern). To address whether the lack of early neural tracking of meter relates to the perception of the meter itself, or to an immature ability to track slower tempi, the current study exposed full-term newborns and young adults to the same rhythm at two different tempi. We hypothesized that in newborns, the patterns of neural responses would be better explained by the tempo of the rhythm (affecting the actual frequencies of the beat, duple and triple periodicities) rather than the metrical structure itself, whereas in adults, the perception of the metrical hierarchy would play a larger role than tempo in orchestrating the pattern of neural responses to the rhythm.

## Materials and Methods

### Participants

High-resolution EEG recordings were obtained from 20 adults (mean age = 21.9 ± 3.63 years old, 9 females) and 23 healthy full-term newborns (mean age = 39.37 ± 1.46 wGA, 10 females). Adults watched a silent movie (Shrek) during the EEG recording in the Auditory Development Lab, McMaster University (Hamilton, Canada). Newborns’ recordings were carried out during sleep at the Department of Obstetrics and Gynecology, Amiens University Hospital (Amiens, France).

**Figure 1.**
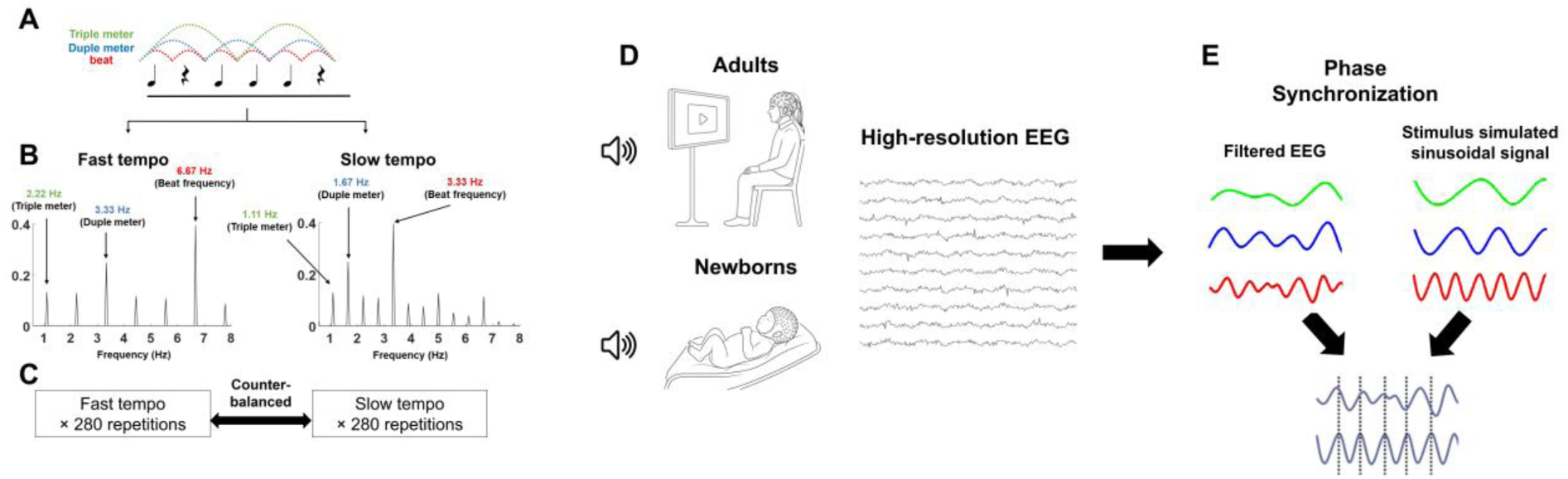
(A) The rhythmic stimulus pattern for the two tempi (inter-beat intervals of 150 ms and 300 ms for fast and slow tempi, respectively). (B) The frequency spectra of fast (left) and slow (right) stimulus sound envelopes. (C) The fast and slow tempo blocks each contained 280 repetitions of the rhythmic pattern, played in counter-balanced order across participants. (D) High-resolution EEG recorded from adults and newborns exposed to the sound stimuli. (E) Brain-stimulus synchronization was calculated for the beat and metrical levels for each tempo separately.

All neonates had appropriate birth weight, size, and head circumference for their term age, an APGAR score > 6 at 5 min, and normal auditory and clinical neurological assessments. Adult participants and one or both parents of infant participants were informed about the study and provided their written informed consent. The McMaster Research Ethics Board at McMaster University (MREB 2164) and the regional ethics committee CPP Ouest I (ID-RCB: 2019-A01534–53) approved the study.

### Auditory stimuli and the experimental paradigm

The auditory stimuli consisted of a 6-beat repeating rhythm in the pattern tone-rest-tone-tone-tone-rest (Phillips-Silver & Trainor, 2005) (Fig. 1A). It was created in two tempi, called hereafter slow and fast tempi. As neural tracking of the musical beat is stronger for low-pitched sounds (Lenc et al., 2018), we chose a pure tone carrier frequency of 130 Hz. The interval between every two successive beats was 300 ms (10-ms rise and 50-ms fall time for tones; corresponding to a beat frequency of 3.33 Hz) for the slow tempo, and 150 ms (5-ms rise and 25-ms fall time; corresponding to a beat frequency of 6.67 Hz) for the fast tempo. The duty-cycle (proportion of the beat during which the tone sounded) was equal to 50% for both fast and slow tempo sequences. Previous research showed that this rhythm can induce the perception of a meter based on grouping by two beats (duple meter; 1.67 Hz for the slow tempo and 3.33 Hz for the fast tempo) or by three beats (triple meter; 1.11 Hz for slow and 2.22 Hz for fast) (Cirelli et al., 2016; Phillips-Silver & Trainor, 2005). The stimuli were created using the open-source software Audacity 2.2.2 and exported as WAV files (https://doi.org/10.17605/OSF.IO/QZHXU).

The magnitude spectra of the acoustic energy in the stimuli corresponding to the slow and fast tempo rhythmic patterns were analyzed by transforming the temporal envelopes of the sounds using a discrete Fourier transform (Fig. 1B). The envelopes of the slow and fast tempo rhythms contained distinct spectral components corresponding to the period of the entire rhythm, its harmonics, and with clear peaks at the beat and metrical levels. The rhythms were designed such that each hierarchical level had the same magnitude of acoustic energy across the slow and fast tempi.

The slow- and fast-tempo rhythmic patterns were presented to participants in two separate blocks, in a randomized order across participants, with each block containing 280 repetitions of the rhythmic pattern (Fig. 1C). The durations of the slow and fast tempo blocks were 504 s (1.8 s × 280 repetitions), and 252 s (0.9 s × 280 repetitions), respectively. The blocks were separated by jittered 3–5 s of silence. The stimuli were delivered through a speaker at 65 dB SPL using Psychtoolbox for MATLAB (Kleiner et al., 2007), positioned at the feet of the newborns and ∼1 meter in front of the adults (Fig. 1D). The total duration of the recording was ∼13 minutes.

### EEG acquisition and preprocessing

EEG signals were collected using HydroCel Geodesic Sensor Nets with 124 channels for newborns and 128 channels for adults using an Electrical Geodesic NetAmps 200 amplifier and Electrical Geodesics Net Station software (version 5). The EEG signals were digitized at a sampling rate of 1000 Hz, with the Cz vertex electrode serving as the reference. Data analyses were performed in MATLAB (MathWorks) using EEGLAB (Delorme & Makeig, 2004), Fieldtrip (Oostenveld et al., 2011), and custom-written MATLAB scripts. EEG data were band-pass filtered between 0.5 – 40 Hz using a two-pass finite impulse response (FIR) filter implemented in EEGLAB (order = 3 cycles of the low-frequency cut-off). A notch filter was applied to attenuate line noise (50 Hz for newborns’ data and 60 Hz for adults’ data). The data were then resampled to 500 Hz. In newborns, the electrodes belonging to the outer ring were removed due to a low signal-to-noise ratio. To have the same number of electrodes across adults and newborns, we also removed the corresponding electrodes from the adults’ data. Following visual inspection, electrodes with artifacts were excluded from further analysis. Eye-blink, eye-movements, and muscle activity in adults’ data, and electrocardiogram artifacts in newborns’ data were removed using independent component analysis implemented in the EEGLAB toolbox. The data were further denoised in EEGLAB using the Artifact Subspace Reconstruction (ASR) algorithm (Plechawska-Wojcik et al., 2018), with a standard deviation cut-off of 10 for both adults and newborns. After re-referencing data to the average reference, electrodes that had been initially excluded were interpolated by replacement with the average of the preprocessed neighboring electrodes. Three newborn participants were excluded from further processing due to a large number of bad electrodes in the frontocentral region resulting from a technical issue related to the EEG cap. Each trial was defined as seven repetitions of the rhythmic pattern, resulting in trial durations of 12.6 s for the slow tempo and 6.3 s for the fast tempo.

### Brain-stimulus synchronization

To evaluate the brain synchronization to the rhythmic sequences at beat and metrical levels, we computed the phase difference between narrow-band filtered (zero-phase FIR, bandwidth 0.5 Hz around the beat, duple, and triple meter frequencies) neural oscillations and the corresponding periodic dynamics of the stimuli (Saadatmehr et al., 2025). To this end, we approximated beat, duple, and triple periodic dynamics by sinusoidal oscillations (Fig. 1E). Each cycle of the sinusoid represented the interval between successive beats or metrical events within the rhythmic sequence. Hilbert transform was then applied to both the filtered EEG signals and the corresponding sinusoidal signals to extract phase time series. The first and last repetition of the rhythmic pattern of each trial were discarded to avoid edge effects. The synchronization index (SI) was computed as the strength of the average difference between the phase time series of EEG data and sinusoidal oscillations across time samples and trials at each electrode for each participant (Moghimi et al., 2020; Staresina et al., 2015):

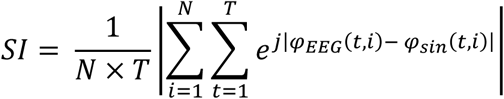

Where *T* represents the number of samples, *N* represents the number of trials, *φ_EEG_*(*t*, *i*) and *φ_sin_*(*t*, *i*) represent the phase values of the filtered EEG signal and the corresponding sinusoidal time series, respectively, at trial *i* and sample *t*.

### Statistical analyses

Statistical analyses were performed in MATLAB (MathWorks) using custom-written MATLAB functions. To assess the SI computed between neural oscillations and periodicities related to beat and metrical levels, surrogate data were generated by randomly permuting the temporal order of the sinusoidal signals independently for each trial (1000 iterations), thereby preserving the amplitude distribution while disrupting their temporal structure. For each shuffle, SI was computed for the shuffled sinusoidal signal and the narrow-band filtered EEG signal at each frequency of interest and each electrode. These SI values were used to create the chance-level distribution of SI across participants. The observed SI values were then compared with the surrogate distribution at each electrode and each frequency of interest using the complementary error function method (Theiler et al., 1992), allowing identification of electrodes where SI was significantly above the chance-level (p < 0.05). To correct for multiple comparisons, we applied the cluster-based permutation correction method with 1000 permutations to find the significant clusters across electrodes (Maris & Oostenveld, 2007). The initial threshold for cluster definition and the minimum number of neighbors were set to p < 0.05 and two, respectively. We reported the cluster-level statistics as the sum of the z-scores (z-scored by permutated SIs) of the electrodes within each permutation-derived cluster, along with the corresponding cluster-level p-value.

To examine whether the synchronization pattern across meters was different across groups, we performed a 2×3×2 mixed-design ANOVA, with group (adults and newborns) as a between-subjects factor and metrical level (beat, duple, and triple meter frequencies) and tempo (slow and fast) as within-subject factors. We averaged SIs across all electrodes to evaluate the brain response at different metrical levels and to remove electrode bias. To assess differences in synchronization patterns between tempi, we also performed a 2×3 repeated-measures ANOVA with metrical level (beat, duple, and triple meter frequencies) and tempo (slow and fast) as within-subject factors separately for each group. The meter x tempo interaction indicated whether the pattern of metrical hierarchy perception was different across slow and fast tempi, separately for adults and newborns. To evaluate the strength of evidence for each effect, we calculated Bayes Factor (BF) using the bayesFactor toolbox in MATLAB (https://github.com/klabhub/bayesFactor). BFs for specific effects were computed as the ratio of BF of the model with the effect of interest to that of the model without that effect.

To quantify the proportion of variance in neural synchronization separately for infants and adults that could be attributed to metrical level (including beat, duple, and triple levels) versus frequency variations within each metrical level, we performed regression-based variance partitioning. Because stimulus frequencies differed across the tempi for each metrical level (triple: 1.11, 2.22 Hz; duple: 1.67, 3.33 Hz; beat: 3.33, 6.67 Hz for slow and fast tempo, respectively), frequency was modeled as nested within metrical levels (3.33 Hz was common between the duple meter frequency of fast tempo and the beat frequency of slow tempo, but they were coded differently in the model). Hierarchical linear regression models were constructed with SI as the dependent variable, including (i) a baseline model containing subject identity to account for between-subject variance, (ii) a model including subject identity and metrical level, and (iii) a model including subject identity and frequency nested within metrical level. The proportion of within-subject variance explained by each factor (i.e., metrical level; frequency within metrical level) was quantified as the change in explained variance (ΔR^2^) when that factor was added to the baseline model containing only subject identity, divided by the total within-subject variance (1-R^2^). As the frequency was nested within the metrical levels, these predictors were orthogonal, which ensured that the variance components were additive and directly comparable (Maxwell et al., 2017).

## Results

### Neural synchronization to the metric hierarchy of slow and fast tempi for adults and newborns

To investigate the synchronization between the EEG neural activity and the rhythmic stimuli, we quantified brain-stimulus synchronization as the strength of phase synchronization (SI) between the simulated sinusoidal signal at the frequency of the metrical levels (beat, duple, and triple) and the corresponding narrow-band filtered neural activity. The topographical scalp distributions of grand-average SI values across participants at these meter-related frequencies (beat, duple, and triple) for adults and newborns for both slow and fast tempi are shown in Fig. 2A and 2B. Each SI value was then compared with the chance level of its corresponding surrogate distribution at each electrode and each frequency of interest (beat, duple, and triple). The cluster-based permutation correction revealed statistically significant clusters in adults at the beat frequency (3.33 Hz, frontoposterior cluster: z = 270.473, p < 0.001 for slow tempo; 6.67 Hz, frontal cluster: z = 115.438, p < 0.001, posterior cluster: z = 23.724, p = 0.024 for fast tempo), duple meter frequency (1.67 Hz, frontoposterior cluster: z = 435.137, p < 0.001 for slow tempo; 3.33 Hz, frontoposterior cluster: z = 260.554, p < 0.001 for fast tempo), and triple meter frequency (1.11 Hz, frontoposterior cluster: z = 249.206, p < 0.001 for slow tempo; 2.22 Hz, frontoposterior cluster: z = 147.75, p < 0.001 for fast tempo).

In newborns, significant clusters were similarly identified across all metrical levels. At the beat frequency, significant SI was observed at 3.33 Hz for the slow tempo (frontoposterior cluster: z = 350.021, p < 0.001) and at 6.67 Hz for the fast tempo (frontal cluster: z = 27, p = 0.008; posterior cluster: z = 26.139, p = 0.011). At the duple meter frequency, significant clusters emerged at 1.67 Hz for the slow tempo (frontal cluster: z = 24.679, p = 0.021; posterior cluster: z = 47.656, p = 0.003) and at 3.33 Hz for the fast tempo (frontal cluster: z = 36.411, p = 0.003; posterior cluster: z = 25.979, p = 0.015). Finally, at the triple meter frequency, significant clusters were found at 1.11 Hz for the slow tempo (posterolateral cluster: z = 25.316, p = 0.025) and at 2.22 Hz for the fast tempo (frontal cluster: z = 17.965, p = 0.046).

### The role of tempo and metrical level in explaining neural synchronization

We further evaluated the pattern of average neural SI across metrical levels for adults and newborns (Fig. 2C and 2D). Visual investigation of Figures 2C and 2D showed that in adults, the pattern of SI variation at the two tempi was relatively the same, although SI tended to peak more dominantly at the duple frequency (1.67 Hz) for the slow tempo than fast tempo. However, in newborns, the SI patterns over different metrical levels showed very different patterns between the two tempi. The mixed-design ANOVA with group as a between-subjects factor and metrical level and tempo as within-subject factors on the SI values revealed a significant three-way interaction (F(2,76) = 5.038, p = 0.009, 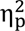 = 0.117, BF_10_ = 8.139), indicating that the modulatory effect of tempo on neural synchronization across metrical levels differed significantly between adults and newborns. Follow-up repeated measures ANOVAs within each group showed that adults exhibited a significant main effect of tempo (F(1,19) = 10.929, p = 0.004, 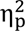 = 0.365, BF_10_ = 8.7), a significant main effect of metrical level (F(2,38) = 8.77, p < 0.001, 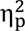 = 0.316, BF_10_ = 27.049) but no tempo × meter interaction (F(2,38) = 1.367, p = 0.267, 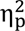 = 0.067, BF_10_ = 0.242), indicating no significant impact of tempo in the pattern of response across the metrical hierarchy. On the other hand, newborns showed no significant main effect of tempo (F(1,19) = 3.675, p = 0.07, 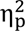 = 0.162, BF_10_ = 0.602) but a significant main effect of metrical level (F(2,38) = 8.77, p = 0.005, 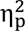 = 0.25, BF_10_ = 12.232) and a significant tempo × meter interaction (F(2,38) = 7.239, p = 0.002, 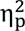 = 0.276, BF_10_ = 46.705). The significant interaction shows that there was a reliable difference in the pattern of SI over different hierarchical levels between fast and slow tempi. Put together, these results suggest that while adults’ neural tracking of the metrical level was not significantly different across tempi, newborns’ neural tracking patterns of metrical level changed from the slow tempo to the fast tempo.

**Figure 2.**
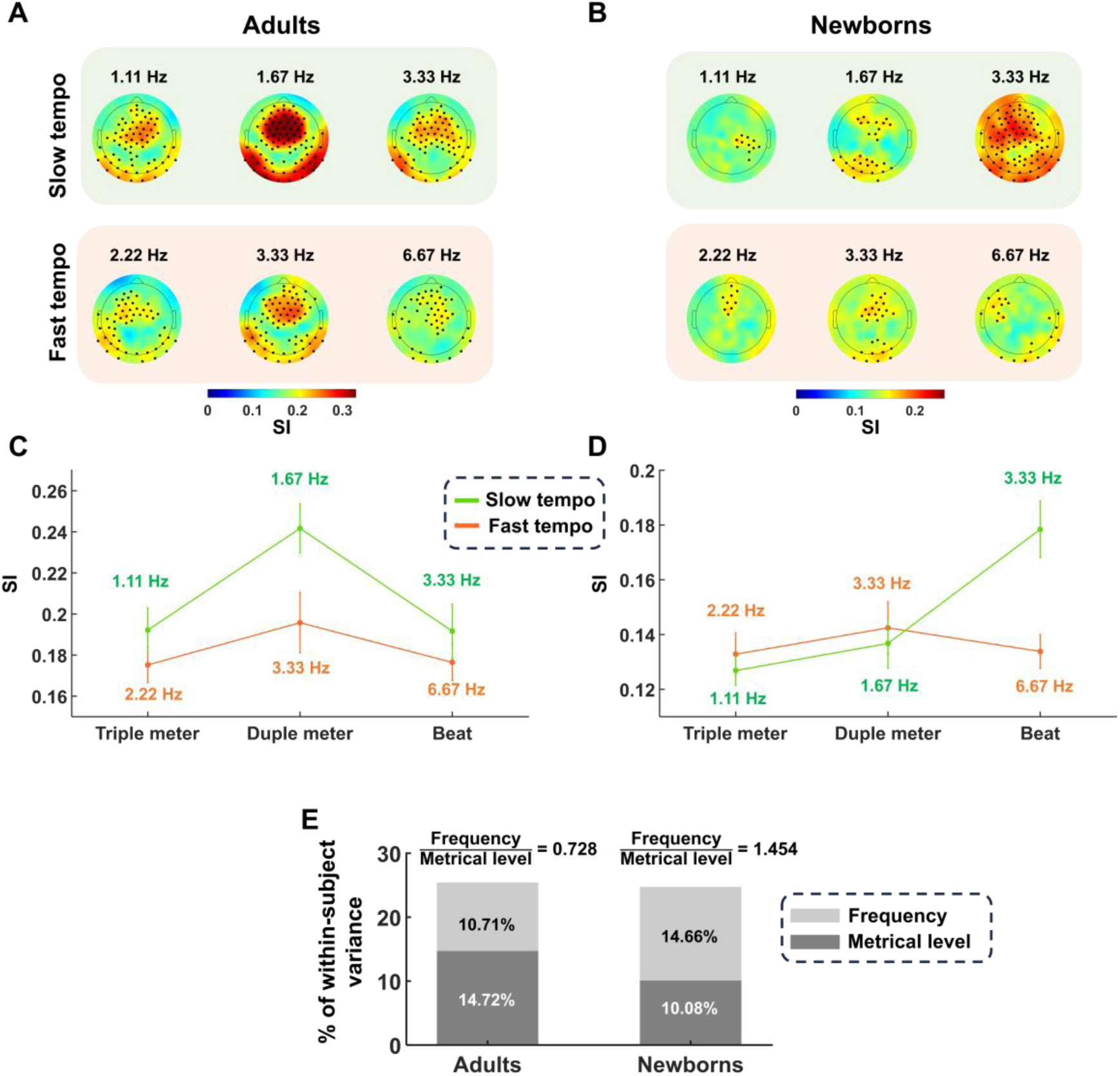
Neural synchronization to rhythmic hierarchy over slow and fast tempi. The topographical scalp distributions for the grand-average SI across metrical levels for adults (A) and newborns (B) at slow (top) and fast (bottom) tempi. The bold dots overlaid on the topographical maps represent significant electrodes after cluster-based permutation correction. C, D, Averaged SI values across all channels for slow and fast tempi in adults and newborns, respectively. (E) The proportion of explained variance by metrical levels and frequency nested within metrical levels for adults and newborns. The role of frequency nested within metrical levels in adults was 0.728 times the role of metrical levels, while this ratio for newborns was 1.454.

To further investigate the role of metrical level and tempo on the neural response to rhythm in adults and newborns, we conducted the regression-based variance partitioning. This analysis in adults showed that metrical level explained 14.72% of within-subject variance (ΔR^2^ = 0.103, F(2,38) = 3.15, p < 0.001), while frequency nested within metrical level accounted for an additional 10.71% (ΔR^2^ = 0.075, F(3,57) = 2.645, p < 0.001). In newborns, metrical level explained 10.08% of within-subject variance (ΔR^2^ = 0.0664, F(2,38) = 3.209, p < 0.001), whereas frequency nested within metrical level explained 14.66% (ΔR^2^ = 0.097, F(3,57) = 3.431, p < 0.001). These results indicated that the total proportion (metrical level + tempo) of within-subject variance explained by metrical level and frequency nested within metrical levels was relatively similar; 25.43% for adults and 24.74% for newborns (Fig. 2E). Next, to more precisely compare the role of each factor between the two groups, we computed the proportion of variance explained by frequency nested within metrical level and metrical level. This proportion was 0.728 for adults and 1.4544 for newborns, demonstrating that frequency at each metrical level played a relatively stronger role than the metrical level in explaining variance in neural responses in newborns compared to adults.

## Discussion

Both music and language rely on hierarchically organized temporal structures that require the integration of information across multiple timescales, including phonemes, syllables, and words in speech, and beat and beat groupings that form the metrical hierarchy in music (Fiveash et al., 2021; Goswami, 2022; Patel, 2003). By presenting a repeating 6-beat rhythm pattern at two tempi (one double that of the other, but with matched duty cycles), containing energy at beat, duple, and triple frequency levels of the metrical hierarchy, we were able to compare the relative contributions of basic frequency (periodicity) and the metrical hierarchy in newborns’ and adults’ neural tracking of the rhythm. We found that for adults, regardless of whether the beat frequency (i.e., tempo) was slower (3.33 Hz) or faster (6.67 Hz), neural responses were closely tied to the metrical hierarchy, with the EEG maximum amplitude response at the duple level of the metrical hierarchy for both tempi (i.e., at 1.67 and 3.33 Hz for the slower and faster tempi, respectively). In contrast, for infants, absolute frequency affected the neural response more than the metrical hierarchy, with the EEG maximum amplitude response at 3.33 Hz regardless of whether it was at the beat (as it was for the slower tempo) or the duple (as it was for the faster tempo) frequency.

While the ability to track slower frequencies appears to develop during the last prenatal trimester (Saadatmehr et al., 2025), it should be noted that the full-term newborns in the present study showed significant neural tracking of all frequencies of interest regardless of tempo. This finding suggests that the newborn-adult differences do not simply reflect an inability of newborns to track slower frequencies, but rather reflect differences in encoding the pattern of responses across the metrical hierarchy of frequencies. Further, infants’ responses to the metrical hierarchy looked most like those of adults at the faster tempo, suggesting that infants might show adult-like responses to the metrical hierarchy, but only over a narrow tempo range. Previous research has demonstrated that also for adults, tempo modulates both behavioral responses and neural following of auditory regularities (Assaneo & Poeppel, 2018; Damsma et al., 2025; Nozaradan et al., 2012; Weineck et al., 2022), but to a much lesser extent that found for the infants in the present study. It is possible, then, that the range of frequencies over which neural tracking is possible expands greatly during the prenatal period, whereas the range over which temporal regularities are integrated over different time scales (i.e., metrical hierarchies) expands postnatally.

Despite extensive investigation of the mechanisms by which adults process metrical hierarchies in both speech and music (Arnal et al., 2015; Kotz et al., 2018; Nozaradan et al., 2011, 2012; Poeppel & Assaneo, 2020; Zalta et al., 2024), their respective roles during the earliest stages of development have remained unclear. Different musical systems preferentially employ different beat groupings or metrical structures (Jacoby & McDermott, 2017) to which infants must become sensitive in order to acquire the musical system in their environment (Hannon & Trainor, 2007; K. Nave et al., 2024). As a consequence, the perception of metrical structure is strongly influenced by long-term experience and learning over the first year (Hannon & Trehub, 2005a, 2005b), while remaining malleable by short-term brief exposure or priming (Flaten et al., 2022). Thus, it makes sense, as our data indicates, that the developing brain initially relies strongly on basic temporal tracking to encode rhythmic input, whereas experience-dependent sensitivity to higher-order metrical structure and top-down influences on rhythm perception emerge later. More broadly, this developmental shift provides insight into how the brain acquires the capacity to represent and integrate information across multiple temporal scales, and how it is constrained by signals that unfold at rates that may be too slow or too rapid for developing neural systems to efficiently process.

From a functional perspective, these findings suggest that early in development, encoding of the auditory environment may preferentially support temporal groupings at intermediate or faster tempo ranges. For example, slower temporal structures corresponding to the grouping of multiple syllables or short words in speech, or longer phrases in music, may be less efficiently encoded than faster groupings such as individual syllables or musical notes. At the same time, more rapid temporal fluctuations may also fall outside the optimal processing range of the newborn auditory system, but this has yet to be tested. Although the optimal frequency observed in the present study (∼3.33 Hz) is somewhat faster than the dominant modulation rates typically reported in infant-directed speech and music (approximately 2 Hz; Goswami, 2022), the current experimental design did not allow a systematic characterization of the optimal temporal tuning range during early development, as it was only possible to test two tempi. The question of the optimal tempo range for periodicity coding and how it changes during the third trimester of gestation was also raised by Saadatmehr et al. (2025) and requires further investigation.

Both endogenous and exogenous factors likely contribute to shaping neural responses to auditory rhythm across development. From a mechanistic perspective, intrinsic maturational changes in neural circuitry—including progressive refinement of excitatory–inhibitory balance (Chini et al., 2022), synaptic pruning (Kostović et al., 2019), and the emergence of faster and more stable oscillatory activity (Bitzenhofer et al., 2020)—may alter the intrinsic temporal dynamics of neural populations (Cross et al., 2025; Wilkinson et al., 2024). Such maturational processes can modify the *eigenfrequency* of large-scale neural networks, thereby influencing the range of external rhythmic frequencies to which the system most efficiently entrains (Zoefel & Kösem, 2024). This framework can account for developmental (Attaheri et al., 2022) and inter-individual differences (in adults) in neural encoding of speech and music, suggesting that shifts in endogenous oscillatory properties may lead to maturational patterns and constrain or facilitate sensitivity to particular temporal scales. However, the developmental trajectory of these intrinsic temporal tuning mechanisms remains largely unexplored, particularly during the earliest stages of postnatal life when oscillatory dynamics are rapidly organizing and reorganizing (Wilkinson et al., 2024).

Evidence from adults also highlights the important contribution of sensorimotor networks and auditory–motor interactions in shaping rhythm perception in both music and speech domains (Fujioka et al., 2012; Keitel et al., 2018; Morillon & Baillet, 2017; Zalta et al., 2024). Auditory rhythm processing recruits motor-related cortical and subcortical structures, even during passive listening (Chen et al., 2006; Grahn & McAuley, 2009; Patel & Iversen, 2014), suggesting that the motor system contributes predictive temporal models that facilitate parsing of rhythmic structure (Large et al., 2023; Tichko et al., 2021; Vuust et al., 2022). Importantly, in adults, auditory–motor coupling during speech perception is itself rate-dependent (Assaneo & Poeppel, 2018), demonstrating that tempo modulates the efficiency of sensorimotor integration and predictive timing mechanisms. Interestingly, the tempo in Assaneo & Poeppel (2018) is around the natural syllable rate for speech, hence suggesting the role of experience in tuning neural interaction for optimal stimulus processing.

Although functional involvement of motor-related regions during auditory rhythm processing has been observed even in newborns (Mashhadi et al., 2025), these networks are unlikely to be specialized or efficiently integrated with the auditory system at this early developmental stage. Converging behavioral evidence shows that as infants acquire motor skills, they display progressively improved tempo tracking (Cirelli & Kragness, 2025; Kragness et al., 2022, 2023), increased movement synchronization (Kirschner & Tomasello, 2009; Nguyen et al., 2025), and greater flexibility in adjusting to faster and slower rhythmic patterns (Kirschner & Tomasello, 2009; Yu et al., 2022; Yu & Myowa, 2021). These developmental changes suggest that emerging motor control may provide additional temporal scaffolding that supports the extraction of higher-order rhythmic regularities at slower tempi. The developmental emergence of greater tempo-flexible neural tracking and increasing sensitivity to metrical organization may reflect the gradual strengthening of auditory–motor coupling. This in turn can provide expanding opportunities for sensorimotor entrainment with development. Such coupling may allow the developing brain to move beyond reliance on basic stimulus-driven temporal constraints toward predictive, hierarchical representations of rhythmic structure (Large et al., 2023; Matthews et al., 2026; Vuust et al., 2022). This perspective also proposes that early rhythm encoding is probably more dominantly constrained by endogenous temporal processing limits, whereas experience-dependent calibration through motor interaction progressively enables the integration of multiple nested temporal scales.

An alternative, non-mutually exclusive explanation of our findings is that developmental differences in neural responses to auditory rhythms may partly reflect maturational changes in the morphology and temporal dynamics of auditory evoked potentials. Changes in response latency, amplitude, and temporal integration windows across development (Daneshvarfard et al., 2019; Graven & Browne, 2008; Saadatmehr et al., 2026; Shafer et al., 2015) could modify the overall neural response to the acoustic envelope of auditory rhythms, potentially influencing frequency-domain measures of neural tracking. This interpretation may be particularly relevant for rhythmic stimuli with sharp acoustic envelope edges (Damsma et al., 2025; Doelling et al., 2019). Because early auditory responses undergo substantial developmental reorganization, the relative contribution of evoked responses and oscillatory alignment may differ between newborns and adults. Although the present study cannot rule out contributions from developmentally-changing evoked responses, the current design does not allow dissociation between evoked-response superposition and genuine oscillatory entrainment. Future studies manipulating envelope dynamics and rhythmic predictability, combined with analytical approaches separating transient and sustained neural activity, will be necessary to clarify these mechanisms.

In the present study, adults were recorded while awake whereas newborns were tested during sleep, a common practice in neonatal EEG studies investigating auditory processing (Bianco et al., 2025; Edalati et al., 2023; Panzani et al., 2023). Importantly, sleep during the neonatal and early infancy periods differs substantially from adult sleep in both its organization and functional properties (Antony et al., 2018; Jenni et al., 2004). Moreover, accumulating evidence suggests that neonates retain robust capacities to process auditory information during sleep (Fló et al., 2025), including speech and musical structure, which are often attenuated in adults during sleep (Sifuentes-Ortega et al., 2022). Consequently, recording newborns during sleep in the current study is unlikely to preclude neural encoding of rhythmic structure.

In the current study, we were not able to examine a wide range of tempi due to constraints on recording duration. This limitation prevented us from addressing questions regarding the optimal tempo ranges for neural rhythm tracking of frequency and metrical structure, as well as from performing direct and systematic comparisons between adults and newborns across multiple temporal scales. Future research could address this issue by implementing a between-participant design that distributes tempo conditions across independent cohorts while preserving protocol feasibility in newborn populations. Furthermore, a more comprehensive characterization of developmental trajectories will require longitudinal approaches that track rhythmic processing across early life while considering environmental influences, such as variability in auditory exposure (Bergelson et al., 2023; Brookman et al., 2020), musical experience (Politimou et al., 2018; Steinberg et al., 2021), and caregiver–infant interaction patterns (Alviar et al., 2026). Such designs would allow a more precise dissociation between maturational and experiential contributions to the development of neural rhythm encoding.

## ACKNOWLEDGMENTS

We thank the participants, parents, and families who consented to take part in the study. This work was supported by Fondation pour l’Audition Grant BabyMusic RD-2021-11 and Agence Nationale de la Recherche ANR-22-CE37-0032.

## DATA AVAILABILITY STATEMENT

The data are not publicly available because participants did not provide consent to share their data outside our research consortium on the consent form. MATLAB code and data matrices are available on OSF: https://doi.org/10.17605/OSF.IO/QZHXU

## References

Alviar, C., Jones, W., & Lense, M. (2026). InCHORRRuS: Infant-Directed Communication Highlights and Organizes Repetition and Redundancy Through Rhythmic Structure. Annals of the New York Academy of Sciences, 1556(1), e70147.

Antony, J. W., Piloto, L., Wang, M., Pacheco, P., Norman, K. A., & Paller, K. A. (2018). Sleep spindle refractoriness segregates periods of memory reactivation. Current Biology, 28(11), 1736–1743. e4.

Arnal, L. H., Doelling, K. B., & Poeppel, D. (2015). Delta–Beta Coupled Oscillations Underlie Temporal Prediction Accuracy. Cerebral Cortex, 25(9), 3077–3085. 10.1093/cercor/bhu103

Assaneo, M. F., & Poeppel, D. (2018). The coupling between auditory and motor cortices is rate-restricted: Evidence for an intrinsic speech-motor rhythm. Science Advances, 4(2), eaao3842.

Attaheri, A., Choisdealbha, Á. N., Di Liberto, G. M., Rocha, S., Brusini, P., Mead, N., Olawole-Scott, H., Boutris, P., Gibbon, S., Williams, I., Grey, C., Flanagan, S., & Goswami, U. (2022). Delta- and theta-band cortical tracking and phase-amplitude coupling to sung speech by infants. NeuroImage, 247, 118698. 10.1016/j.neuroimage.2021.118698

Bergelson, E., Soderstrom, M., Schwarz, I.-C., Rowland, C. F., Ramírez-Esparza, N., R. Hamrick, L., Marklund, E., Kalashnikova, M., Guez, A., & Casillas, M. (2023). Everyday language input and production in 1,001 children from six continents. Proceedings of the National Academy of Sciences, 120(52), e2300671120.

Bianco, R., Tóth, B., Bigand, F., Nguyen, T., Sziller, I., Háden, G. P., Winkler, I., & Novembre, G. (2025). Human newborns form musical predictions based on rhythmic but not melodic structure. 10.1101/2025.02.19.639016

Bitzenhofer, S. H., Pöpplau, J. A., & Hanganu-Opatz, I. (2020). Gamma activity accelerates during prefrontal development. eLife, 9, e56795. 10.7554/eLife.56795

Brookman, R., Kalashnikova, M., Conti, J., Rattanasone, N. X., Grant, K.-A., Demuth, K., & Burnham, D. (2020). Depression and anxiety in the postnatal period: An examination of infants’ home language environment, vocalizations, and expressive language abilities. Child Development, 91(6), e1211–e1230.

Chen, J. L., Zatorre, R. J., & Penhune, V. B. (2006). Interactions between auditory and dorsal premotor cortex during synchronization to musical rhythms. Neuroimage, 32(4), 1771–1781.

Chini, M., Pfeffer, T., & Hanganu-Opatz, I. (2022). An increase of inhibition drives the developmental decorrelation of neural activity. Elife, 11, e78811.

Choi, D., Batterink, L. J., Black, A. K., Paller, K. A., & Werker, J. F. (2020). Preverbal Infants Discover Statistical Word Patterns at Similar Rates as Adults: Evidence From Neural Entrainment. Psychological Science, 31(9), 1161–1173. 10.1177/0956797620933237

Cirelli, L. K., & Kragness, H. E. (2025). The Development of Dance in Early Childhood. Current Directions in Psychological Science, 09637214251323490. 10.1177/09637214251323490

Cirelli, L. K., Spinelli, C., Nozaradan, S., & Trainor, L. J. (2016). Measuring Neural Entrainment to Beat and Meter in Infants: Effects of Music Background. Frontiers in Neuroscience, 10. 10.3389/fnins.2016.00229

Cirelli, L. K., Trehub, S. E., & Trainor, L. J. (2018). Rhythm and melody as social signals for infants: Rhythm and melody as social signals for infants. Annals of the New York Academy of Sciences, 1423(1), 66–72. 10.1111/nyas.13580

Cross, Z. R., Gray, S. M., Dede, A. J., Rivera, Y. M., Yin, Q., Vahidi, P., Rau, E. M., Cyr, C., Holubecki, A. M., & Asano, E. (2025). The development of aperiodic neural activity in the human brain. Nature Human Behaviour, 1–16.

Damsma, A., de Roo, M., Doelling, K., Bazin, P.-L., & Bouwer, F. L. (2025). Tempo-dependent selective enhancement of neural responses at the beat frequency can be mimicked by both an oscillator and an evoked model. Cerebral Cortex, 35(9), bhaf258.

Daneshvarfard, F., Abrishami Moghaddam, H., Dehaene-Lambertz, G., Kongolo, G., Wallois, F., & Mahmoudzadeh, M. (2019). Neurodevelopment and asymmetry of auditory-related responses to repetitive syllabic stimuli in preterm neonates based on frequency-domain analysis. Scientific Reports, 9(1), 10654.

Delorme, A., & Makeig, S. (2004). EEGLAB: an open source toolbox for analysis of single-trial EEG dynamics including independent component analysis. Journal of Neuroscience Methods, 134(1), 9–21.

Doelling, K. B., Assaneo, M. F., Bevilacqua, D., Pesaran, B., & Poeppel, D. (2019). An oscillator model better predicts cortical entrainment to music. Proceedings of the National Academy of Sciences, 116(20), 10113–10121.

Edalati, M., Wallois, F., Ghostine, G., Kongolo, G., Trainor, L. J., & Moghimi, S. (2024). Neural oscillations suggest periodicity encoding during auditory beat processing in the premature brain. Developmental Science, 27(6), e13550.

Edalati, M., Wallois, F., Safaie, J., Ghostine, G., Kongolo, G., Trainor, L. J., & Moghimi, S. (2023). Rhythm in the premature neonate brain: Very early processing of auditory beat and meter. *The Journal of Neuroscience*, JN-RM-1100–22. 10.1523/JNEUROSCI.1100-22.2023

Fiveash, A., Bedoin, N., Gordon, R. L., & Tillmann, B. (2021). Processing rhythm in speech and music: Shared mechanisms and implications for developmental speech and language disorders. Neuropsychology, 35(8), 771–791. 10.1037/neu0000766

Flaten, E., Marshall, S. A., Dittrich, A., & Trainor, L. J. (2022). Evidence for top-down metre perception in infancy as shown by primed neural responses to an ambiguous rhythm. European Journal of Neuroscience, 55(8), 2003–2023. 10.1111/ejn.15671

Fló, A., Benjamin, L., Palu, M., & Dehaene-Lambertz, G. (2025). Statistical learning beyond words in human neonates. eLife, 13, RP101802. 10.7554/eLife.101802

Fujioka, T., Trainor, L. J., Large, E. W., & Ross, B. (2012). Internalized Timing of Isochronous Sounds Is Represented in Neuromagnetic Beta Oscillations. The Journal of Neuroscience, 32(5), 1791–1802. 10.1523/JNEUROSCI.4107-11.2012

Goswami, U. (2022). Language acquisition and speech rhythm patterns: An auditory neuroscience perspective. Royal Society Open Science, 9(7).

Grahn, J. A., & McAuley, J. D. (2009). Neural bases of individual differences in beat perception. NeuroImage, 47(4), 1894–1903.

Graven, S. N., & Browne, J. V. (2008). Auditory development in the fetus and infant. Newborn and Infant Nursing Reviews, 8(4), 187–193.

Háden, G. P., Bouwer, F. L., Honing, H., & Winkler, I. (2024). Beat processing in newborn infants cannot be explained by statistical learning based on transition probabilities. Cognition, 243, 105670. 10.1016/j.cognition.2023.105670

Háden, G. P., Honing, H., Török, M., & Winkler, I. (2015). Detecting the temporal structure of sound sequences in newborn infants. International Journal of Psychophysiology, 96(1), 23–28. 10.1016/j.ijpsycho.2015.02.024

Hannon, E. E., & Trainor, L. J. (2007). Music acquisition: Effects of enculturation and formal training on development. Trends in Cognitive Sciences, 11(11), 466–472. 10.1016/j.tics.2007.08.008

Hannon, E. E., & Trehub, S. E. (2005a). Metrical Categories in Infancy and Adulthood. Psychological Science, 16(1), 48–55. 10.1111/j.0956-7976.2005.00779.x

Hannon, E. E., & Trehub, S. E. (2005b). Tuning in to musical rhythms: Infants learn more readily than adults. Proceedings of the National Academy of Sciences, 102(35), 12639–12643. 10.1073/pnas.0504254102

Jacoby, N., & McDermott, J. H. (2017). Integer ratio priors on musical rhythm revealed cross-culturally by iterated reproduction. Current Biology, 27(3), 359–370.

Jenni, O. G., Borbély, A. A., & Achermann, P. (2004). Development of the nocturnal sleep electroencephalogram in human infants. *American Journal of Physiology-Regulatory*, Integrative and Comparative Physiology, 286(3), R528–R538.

Keitel, A., Gross, J., & Kayser, C. (2018). Perceptually relevant speech tracking in auditory and motor cortex reflects distinct linguistic features. PLoS Biology, 16(3), e2004473.

Kirschner, S., & Tomasello, M. (2009). Joint drumming: Social context facilitates synchronization in preschool children. Journal of Experimental Child Psychology, 102(3), 299–314. 10.1016/j.jecp.2008.07.005

Kleiner, M., Brainard, D., & Pelli, D. (2007). What’s new in Psychtoolbox*-*3*?*

Kostović, I., Sedmak, G., & Judaš, M. (2019). Neural histology and neurogenesis of the human fetal and infant brain. Neuroimage, 188, 743–773.

Kotz, S. A., Ravignani, A., & Fitch, W. T. (2018). The Evolution of Rhythm Processing. Trends in Cognitive Sciences, 22(10), 896–910. 10.1016/j.tics.2018.08.002

Kragness, H. E., Anderson, L., Chow, E., Schmuckler, M., & Cirelli, L. K. (2023). Musical groove shapes children’s free dancing. Developmental Science, 26(1), e13249. 10.1111/desc.13249

Kragness, H. E., Ullah, F., Chan, E., Moses, R., & Cirelli, L. K. (2022). Tiny dancers: Effects of musical familiarity and tempo on children’s free dancing. Developmental Psychology, 58, 1277–1285. 10.1037/dev0001363

Kuhl, P. K. (2011). Social Mechanisms in Early Language Acquisition: Understanding Integrated Brain Systems Supporting Language. Oxford University Press. 10.1093/oxfordhb/9780195342161.013.0043

Large, E. W., Roman, I., Kim, J. C., Cannon, J., Pazdera, J. K., Trainor, L. J., Rinzel, J., & Bose, A. (2023). Dynamic models for musical rhythm perception and coordination. Frontiers in Computational Neuroscience, 17, 1151895. 10.3389/fncom.2023.1151895

Lenc, T., Keller, P. E., Varlet, M., & Nozaradan, S. (2018). Neural tracking of the musical beat is enhanced by low-frequency sounds. Proceedings of the National Academy of Sciences, 115(32), 8221–8226. 10.1073/pnas.1801421115

Lenc, T., Peter, V., Hooper, C., Keller, P. E., Burnham, D., & Nozaradan, S. (2023). Infants show enhanced neural responses to musical meter frequencies beyond low-level features. Developmental Science, 26(5), e13353. 10.1111/desc.13353

Lense, M. D., Ladányi, E., Rabinowitch, T.-C., Trainor, L., & Gordon, R. (2021). Rhythm and timing as vulnerabilities in neurodevelopmental disorders. Philosophical Transactions of the Royal Society B: Biological Sciences, 376(1835), 20200327. 10.1098/rstb.2020.0327

Maris, E., & Oostenveld, R. (2007). Nonparametric statistical testing of EEG-and MEG-data. Journal of Neuroscience Methods, 164(1), 177–190.

Mashhadi, A. R., Wallois, F., Edalati, M., Levé, F., Stamatiadis, A., Chazal, C., Trainor, L., & Moghimi, S. (2025). Neural encoding of auditory rhythm beyond cortical auditory areas before the age of term. Iscience, 28(9).

Matthews, T. E., Vuust, P., & Cannon, J. (2026). An Active Inference Model of Meter Perception and the Urge to Move to Music. Annals of the New York Academy of Sciences, 1556(1), e70129.

Maxwell, S. E., Delaney, H. D., & Kelley, K. (2017). Designing experiments and analyzing data: A model comparison perspective. Routledge.

Moghimi, S., Shadkam, A., Mahmoudzadeh, M., Calipe, O., Panzani, M., Edalati, M., Ghorbani, M., Routier, L., & Wallois, F. (2020). The intimate relationship between coalescent generators in very premature human newborn brains: Quantifying the coupling of nested endogenous oscillations. Human Brain Mapping, 41(16), 4691–4703. 10.1002/hbm.25150

Morillon, B., & Baillet, S. (2017). Motor origin of temporal predictions in auditory attention. Proceedings of the National Academy of Sciences, 114(42). 10.1073/pnas.1705373114

Nave, K., Carrillo, C., Jacoby, N., Trainor, L., & Hannon, E. (2024). The development of rhythmic categories as revealed through an iterative production task. Cognition, 242, 105634.

Nave, K. M., Carrillo, C., Jacoby, N., Trainor, L., & Hannon, E. (2021). *The Development of Rhythmic Categories as Revealed Through an Iterative Production Task* [Preprint]. PsyArXiv. 10.31234/osf.io/wvb9k

Nave-Blodgett, J. E., Snyder, J. S., & Hannon, E. E. (2021a). Auditory superiority for perceiving the beat level but not measure level in music. Journal of Experimental Psychology: Human Perception and Performance, 47(11), 1516.

Nave-Blodgett, J. E., Snyder, J. S., & Hannon, E. E. (2021b). Hierarchical beat perception develops throughout childhood and adolescence and is enhanced in those with musical training. Journal of Experimental Psychology: General, 150(2), 314–339. 10.1037/xge0000903

Nguyen, T., Bigand, F., Reisner, S., Koul, A., Bianco, R., Markova, G., Hoehl, S., & Novembre, G. (2025). Development of auditory and spontaneous movement responses to music over the first year of life. bioRxiv, 2025.05. 04.649695.

Nozaradan, S., Peretz, I., Missal, M., & Mouraux, A. (2011). Tagging the neuronal entrainment to beat and meter. Journal of Neuroscience, 31(28), 10234–10240.

Nozaradan, S., Peretz, I., & Mouraux, A. (2012). Selective neuronal entrainment to the beat and meter embedded in a musical rhythm. Journal of Neuroscience, 32(49), 17572–17581.

Oostenveld, R., Fries, P., Maris, E., & Schoffelen, J.-M. (2011). FieldTrip: Open source software for advanced analysis of MEG, EEG, and invasive electrophysiological data. Computational Intelligence and Neuroscience, 2011, 1–9.

Panzani, M., Mahmoudzadeh, M., Wallois, F., & Dehaene-Lambertz, G. (2023). Detection of regularities in auditory sequences before and at term-age in human neonates. NeuroImage, 284, 120428.

Patel, A. D. (2003). Language, music, syntax and the brain. Nature Neuroscience, 6(7), 674–681. 10.1038/nn1082

Patel, A. D., & Iversen, J. R. (2014). The evolutionary neuroscience of musical beat perception: The Action Simulation for Auditory Prediction (ASAP) hypothesis. Frontiers in Systems Neuroscience, 8. 10.3389/fnsys.2014.00057

Phillips-Silver, J., & Trainor, L. J. (2005). Feeling the Beat: Movement Influences Infant Rhythm Perception. Science, 308(5727), 1430–1430. 10.1126/science.1110922

Plechawska-Wojcik, M., Kaczorowska, M., & Zapala, D. (2018). The artifact subspace reconstruction (ASR) for EEG signal correction. A comparative study. International Conference on Information Systems Architecture and Technology, 125–135.

Poeppel, D., & Assaneo, M. F. (2020). Speech rhythms and their neural foundations. Nature Reviews Neuroscience, 21(6), 322–334. 10.1038/s41583-020-0304-4

Politimou, N., Stewart, L., Müllensiefen, D., & Franco, F. (2018). Music@Home: A novel instrument to assess the home musical environment in the early years. PLOS ONE, 13(4), e0193819. 10.1371/journal.pone.0193819

Saadatmehr, B., Edalati, M., Wallois, F., Ghostine, G., Kongolo, G., Flaten, E., Tillmann, B., Trainor, L., & Moghimi, S. (2025). Auditory rhythm encoding during the last trimester of human gestation: From tracking the basic beat to tracking hierarchical nested temporal structures. Journal of Neuroscience, 45(4).

Saadatmehr, B., Gallard, A., Edalati, M., Kongolo, G., Ghostine, G., Chazal, C., Brunel, P., David, O., Wallois, F., & Moghimi, S. (2026). Brain complexity in response to auditory stimulation improves evaluation of cerebral maturation in premature newborns. Pediatric Research, 1–8.

Savage, P. E., Loui, P., Tarr, B., Schachner, A., Glowacki, L., Mithen, S., & Fitch, W. T. (2021). Music as a coevolved system for social bonding. Behavioral and Brain Sciences, 44, e59.

Shafer, V. L., Yan, H. Y., & Wagner, M. (2015). Maturation of cortical auditory evoked potentials (CAEPs) to speech recorded from frontocentral and temporal sites: Three months to eight years of age. International Journal of Psychophysiology, 95(2), 77–93.

Sifuentes-Ortega, R., Lenc, T., Nozaradan, S., & Peigneux, P. (2022). Partially preserved processing of musical rhythms in REM but not in NREM sleep. Cerebral Cortex, 32(7), 1508–1519.

Snyder, J. S., Gordon, R. L., & Hannon, E. E. (2024). Theoretical and empirical advances in understanding musical rhythm, beat and metre. Nature Reviews Psychology, 3(7), 449–462. 10.1038/s44159-024-00315-y

Soley, G., & Hannon, E. E. (2010). Infants prefer the musical meter of their own culture: A cross-cultural comparison. Developmental Psychology, 46(1), 286.

Staresina, B. P., Bergmann, T. O., Bonnefond, M., Van Der Meij, R., Jensen, O., Deuker, L., Elger, C. E., Axmacher, N., & Fell, J. (2015). Hierarchical nesting of slow oscillations, spindles and ripples in the human hippocampus during sleep. Nature Neuroscience, 18(11), 1679–1686.

Steinberg, S., Shivers, C. M., Liu, T., Cirelli, L. K., & Lense, M. D. (2021). Survey of the home music environment of children with various developmental profiles. Journal of Applied Developmental Psychology, 75, 101296.

Theiler, J., Eubank, S., Longtin, A., Galdrikian, B., & Farmer, J. D. (1992). Testing for nonlinearity in time series: The method of surrogate data. Physica D: Nonlinear Phenomena, 58(1–4), 77–94.

Tichko, P., Kim, J. C., & Large, E. W. (2021). Bouncing the network: A dynamical systems model of auditory–vestibular interactions underlying infants’ perception of musical rhythm. Developmental Science, 24(5), e13103.

Tillmann, B., Goswami, U., & Moghimi, S. (2025). Rhythm Processing Across Development: Origins, Links to Language Processing, and Perspectives for Intervention. Annals of the New York Academy of Sciences.

Trainor, L. J., & Marsh-Rollo, S. (2018). Rhythm, meter, and timing: The heartbeat of musical development.

Vuust, P., Heggli, O., A., Friston, K., J., & Kringelbach, M., L. (2022). Music in the brain. Nature Reviews Neroscience, 23(5), 287–305.

Weineck, K., Wen, O. X., & Henry, M. J. (2022). Neural synchronization is strongest to the spectral flux of slow music and depends on familiarity and beat salience. Elife, 11, e75515.

Werker, J. F., & Tees, R. C. (2005). Speech perception as a window for understanding plasticity and commitment in language systems of the brain. Developmental Psychobiology: The Journal of the International Society for Developmental Psychobiology, 46(3), 233–251.

Wilkinson, C. L., Yankowitz, L. D., Chao, J. Y., Gutiérrez, R., Rhoades, J. L., Shinnar, S., Purdon, P. L., & Nelson, C. A. (2024). Developmental trajectories of EEG aperiodic and periodic components in children 2–44 months of age. Nature Communications, 15(1), 5788.

Winkler, I., Háden, G. P., Ladinig, O., Sziller, I., & Honing, H. (2009). Newborn infants detect the beat in music. Proceedings of the National Academy of Sciences, 106(7), 2468–2471.

Yu, L., & Myowa, M. (2021). The early development of tempo adjustment and synchronization 8 during joint drumming: A study of 18- to 42-month-old children. Infancy, 26(4), 635–646. 10.1111/infa.12403

Yu, L., Todoriki, K., & Myowa, M. (2022). From spontaneous rhythmic engagement to joint drumming: A gradual development of flexible coordination at approximately 24 months of age. Frontiers in Psychology, 13, 907834. 10.3389/fpsyg.2022.907834

Zalta, A., Large, E. W., Schön, D., & Morillon, B. (2024). Neural dynamics of predictive timing and motor engagement in music listening. Science Advances.

Zoefel, B., & Kösem, A. (2024). Neural tracking of continuous acoustics: Properties, speech-specificity and open questions. European Journal of Neuroscience, 59(3), 394–414. 10.1111/ejn.16221

